# Estimating parameters of a stochastic cell invasion model with fluorescent cell cycle labelling using Approximate Bayesian Computation

**DOI:** 10.1101/2021.04.20.440712

**Authors:** Michael J Carr, Matthew J Simpson, Christopher Drovandi

**Affiliations:** Queensland University of Technology

**Keywords:** Sequential Monte Carlo, SMC-ABC, Cell proliferation, Cell motility, Random walk model

## Abstract

We develop a parameter estimation method based on approximate Bayesian computation (ABC) for a stochastic cell invasion model using fluorescent cell cycle labeling with proliferation, migration, and crowding effects. Previously, inference has been performed on a deterministic version of the model fitted to cell density data, and not all the parameters were identifiable. Considering the stochastic model allows us to harness more features of experimental data, including cell trajectories and cell count data, which we show overcomes the parameter identifiability problem. We demonstrate that, whilst difficult to collect, cell trajectory data can provide more information about the parameters of the cell invasion model. To handle the intractability of the likelihood function of the stochastic model, we use an efficient ABC algorithm based on sequential Monte Carlo. Rcpp and MATLAB implementations of the simulation model and ABC algorithm used in this study are available at https://github.com/michaelcarr-stats/FUCCI.

## 1 Introduction

Australia and New Zealand have the highest incidence rates of melanoma in the world, followed by northern America and northern Europe (Parkin et al., 2005). In Australia, melanoma is the third most common diagnosed form of cancer (Australian Institute of Health and Welfare, 2018). Since the 1960s, Australia’s primary strategy to reduce overall mortality rates has been targeted at early prevention and detection (Giblin & Thomas, 2007). However, a better understanding of the mechanisms which control cell invasion is necessary in order to improve or establish new treatment measures.

The underlying mechanisms of cell invasion we consider are combined cell proliferation and cell migration. Cell proliferation is a four-stage sequence consisting of gap 1 (G1), synthesis (S), gap 2 (G2), and mitosis (M) where the cell divides into two daughter cells, each of which return to the G1 phase (Haass et al., 2014). Improvements in technology have enabled us to visualise different phases of the cell cycle in real time using Fluorescent Ubiquitination-based Cell Cycle Indicator (FUCCI) technology (Sakaue-Sawano et al., 2008). FUCCI technology involves two fluorescent probes which emit red fluorescence when the cells are in G1 phase and green fluorescence when in S/G2/M phases. During the transition between G1 and S phase, both probes are active (giving the impression that the cell fluoresces yellow), allowing the visualisation of the early S phase, which we refer to as eS. Experiments using FUCCI-transduced melonama cells are becoming increasingly important in cancer research because many drug treatments target different phases of the cell cycle (Haass & Gabrielli, 2017).

The development of simulation models offer us a quick and inexpensive alternative to *in vitro* experiments. In this study, we adopt the cell invasion model of scratch assay experiments developed by Simpson et al. (2018). This model describes a discrete exclusion based random walk on a two-dimensional (2D) hexagonal lattice. Furthermore, this model involves treating the entire population of agents as three subpopulations that correspond to the red, yellow and green phases of the cell cycle as identified by FUCCI. Agents transition through the cell cycle, while simultaneously undergoing a nearest neighbour random walk, with exclusion, to model cell migration. This previous study did not perform any parameter inference or calibrate the model to experimental data. The primary focus of this present work is to apply Bayesian methods to recover parameter estimates for the model and the associated distribution of uncertainly around them. However, standard Bayesian approaches rely on the computation of the likelihood function which is often intractable in complex stochastic models. We overcome this limitation by applying Approximate Bayesian Computation (ABC) methods.

Simpson et al. (2020) investigate parameter identifiability in a deterministic partial differential equation of FUCCI scratch assay experiments. By using a simpler model, their study is able to adopt standard Bayesian approaches to parameter estimation since the likelihood function is tractable. Using a Markov Chain Monte Carlo (MCMC) framework and cell density data, this study found cell diffusivities were practically non-identifiable when they considered the case where cell migration rate depends on the cell cycle phase. Although, this study does not consider other types of data which may be more informative of the underlying mechanisms. Here, we address the limitations Simpson et al. (2020) identify by modelling individual cell behaviour with a stochastic model which allows the generation of numerous data types. Indeed, we take full advantage of the flexibility of the stochastic model in this study and combine multiple data types (the number of cells in each phase and cell trajectory data accounting for different phases) to improve parameter identifiability. However, working with cell trajectory data can be challenging, and these challenges include time consuming effort to manually track cells and the need for the cell density to be low to make cell tracking easier. Models which can avoid using cell trajectory data is an active area of research (Hywood et al., 2021), but we find using the Simpson et al. (2018) model which incorporates cell trajectory data leads to a good outcome.

Many other studies have explored modelling and/or parameter estimation in cell invasion models (Cai et al., 2007; Maini et al., 2004; Savla et al., 2004; Swanson, 2008; Takamizawa et al., 1997; Vo et al., 2015). Notably, Vo et al. (2015) estimate the parameters of a stochastic cell spreading model of an expanding population of fibroblast cells in a 2D circular barrier assay without cell cycle labelling. While ABC methods have previously been considered in stochastic cell spreading models, such as the Vo et al. (2015) study, they have never before been considered with FUCCI models and/or data. Prior to Vo et al. (2015), cell invasion models were usually defined by deterministic partial differential equation and when performing parameter inference, they usually used trial and error based approaches (Takamizawa et al., 1997) or non-linear least squares estimation (Cai et al., 2007; Maini et al., 2004; Savla et al., 2004; Swanson, 2008). However, these approaches to parameter estimation are unable to quantify the uncertainty around the point estimates. Rcpp and MATLAB implementations of the simulation model and ABC algorithm used in this study are available at https://github.com/michaelcarr-stats/FUCCI.

The paper is structured as follows. In Section 2, we introduce the experimental data and the process by which it is collected. Section 3 describes the simulation model, the parameter inference method used, and our prior knowledge on the model parameters. In Section 4 we explain the image analysis process and present the inference results when using synthetic and experimental data sets. Discussion of results, future work and concluding remarks are presented in Section 5.

## 2 Data

2D scratch assay experiments are a good screening tool for more complex experimental models, as they are low cost, allow for easy data interpretation and readily allow control of oxygen, nutrients and drug supply (Beaumont et al., 2014; Santiago-Walker et al., 2009). We adopt data from a study conducted by Vittadello et al. (2018) where a scratch assay is used to examine melanoma cell proliferation and migration in real time with FUCCI technology. The experiment is initialised by placing a small population of cells and a growth medium in a culture dish (Figure 1 (a)) to create a uniform 2D monolayer of cells. Next, a sharp-tipped instrument is used to make a scratch is made in the monolayer of cells (Figure 1 (b)). Finally, the cells are observed at regular intervals as they proliferate and migrate into the newly created gap over the following 48 hours. For this study, we adopt the data from the experiments with WM983C FUCCI-transduced melonoma cells and present still images captured at 0 and 48 hours in Figure 1 (c)-(d), respectively. A major advantage of 2D scratch assay experiments is the multitude of different data types which can be easily recovered. The data types which we explore later include the number of cells in each population, position of cell populations, and cell trajectory data (Figure 1 (e)). It is important to consider the size of the imaged region compared to the culture plate (Figure 1 (b)) because the boundaries of the imaged region are not physical boundaries. Since the cell density outside of the scratched region is approximately uniform, with no macroscopic density gradients away from the leading edge, the net flux of cells across the boundary will be zero (Simpson et al., 2018). Therefore, the appropriate mathematical boundary conditions along the vertical boundaries will be zero net flux.

**Figure 1:**
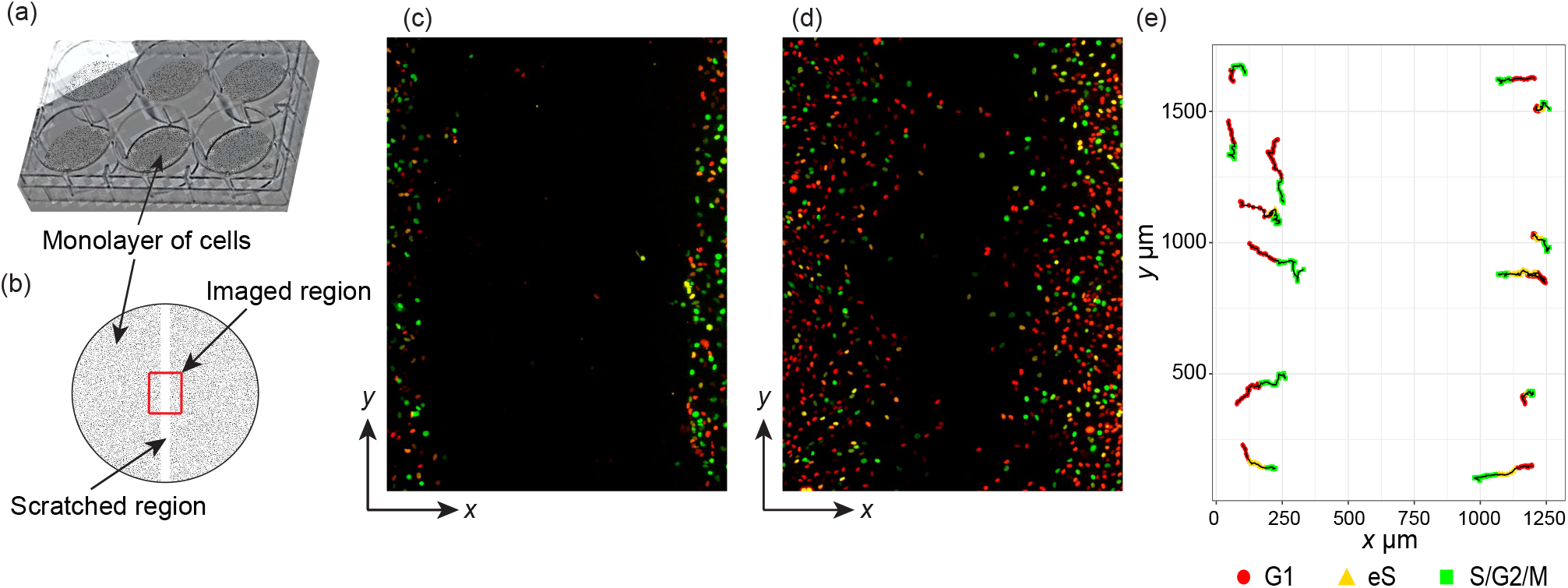
Experimental procedure and data. (a)-(b) Explains the experimental procedure and boundary conditions for simulation models. (a) Photograph of 6 culture plates commonly used with a uniform monolayer of cells. (b) Schematic showing the uniform cell monolayer (shaded), scratched region (white), and imaged region (outlined in red) in a 35 mm culture plate. (c)-(d) Experimental images, both 1309.09 x 1745.35 μm, of WM983C FUCCI-transduced melanoma cells at 0 and 48 hours, respectively. Images reproduced with permission from Vittadello et al. (2018) (e) Cell trajectory data of a select few cells recorded through red to green phases travelling inward to fill scratched region.

## 3 Methods

### 3.1 Simulation model

We adopt the random walk model developed by Simpson et al. (2018) on a 2D hexagonal lattice. Each lattice site has diameter Δ = 20 μm (the average cell diameter (Treloar et al., 2013)) and is associated with a set of unique Cartesian coordinates,

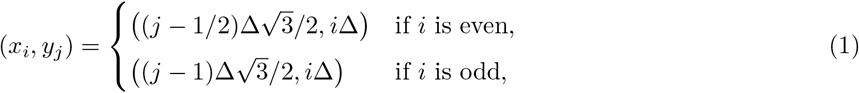

where *i* and *j* are the respective row and column indices. To mimic scratch assay experiments, cells in G1 phase are represented by red agents, cells in eS phase are represented by yellow agents, and cells in S/G2/M phase are represented by green agents. Agents are permitted to transition through phases of the cell cycle and undergo a nearest neighbour random walk by simulating from a Markov process using the Gillespie algorithm (Gillespie, 1977) which is presented in Appendix A.

To simulate cell migration, agents undergo a nearest neighbour random walk at rates *M_r_,M_y_,M_g_* per hour for red, yellow and green agents, respectively (Figure 2 (a)-(f)). Potential movement events involve randomly selecting the target site from the set of six nearest-neighbouring lattice sites, with the movement event being successful only if the target site is vacant. In this way crowding effects are simply accommodated. To simulate transitions through the cell cycle, red agents are allowed to transition into yellow agents at rate *R_r_* per hour (Figure 2 (h)-(i)), yellow agents to green agents at rate *R_y_* per hour (Figure 2 (i)-(j)) and green agents into two red daughter agents at rate *R_g_* per hour (Figure 2 (j)-(k)). While we assume that the red-to-yellow and yellow-to-green transitions are unaffected by crowding, we model crowding effects for the green-to-red transition by aborting transitions where the additional red daughter agent would be placed onto an occupied lattice site. By prohibiting multiple agents from occupying the same lattice site, we are able to realistically incorporate crowding effects (Ermentrout & Edelstein-Keshet, 1993; Johnston et al., 2016).

**Figure 2:**
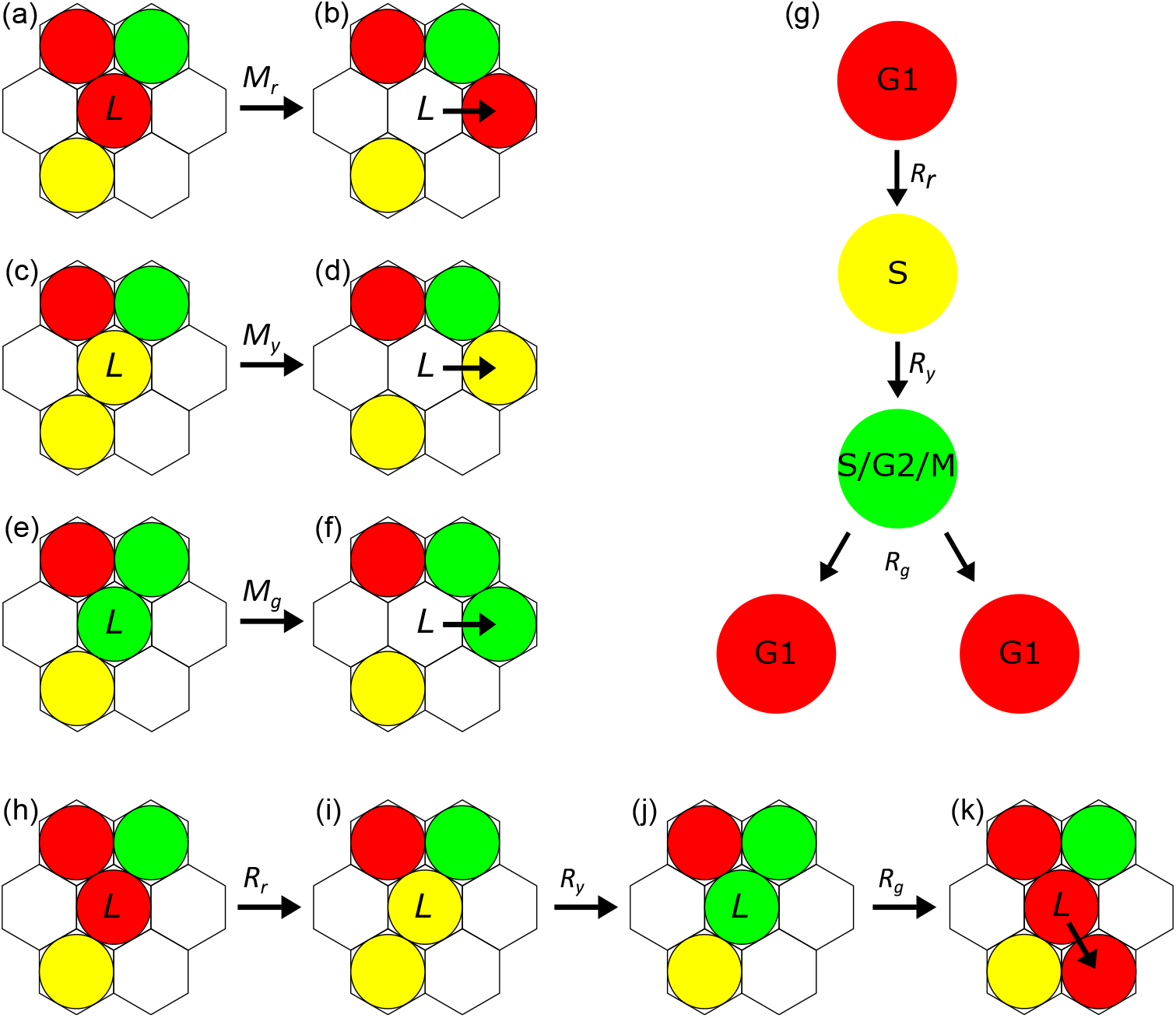
Cell migration and proliferation. (a-f) An agent at lattice site *L* will attempt to migrate to the six neighbouring lattice sites, successfully migrating if the selected site is vacant. (g) Schematic showing the progression through the G1 phase (red), early S phase (yellow) and S/G2/M phase (green) for FUCCI. (h-k) Agent transition through the cell cycle and proliferation. (k) A green agent (S/G2/M phase) at lattice site *L* will successfully divide and transition if the randomly selected neighbouring site is vacant.

The Simpson et al. (2018) model is dependent on the initial geometry, boundary conditions, the lattice spacing Δ, and the cell cycle transition and motility rates. Since we have reasonable estimates for Δ (Treloar et al., 2013) and we calibrate the initial geometry and boundary conditions to the experimental data, our study is concerned with estimating the unknown cell cycle transition and motility parameters. In a Bayesian setting, the unknown model parameters, *θ* = (*R_r_, R_y_, R_g_, M_r_, M_y_, M_g_*), and the uncertainty around them can be quantified by the posterior distribution; which is dependent on the likelihood and the prior distribution. However, while the Markov process model can capture the stochastic nature of cell proliferation and migration, when the dimension of the generator matrix (a matrix of rate parameters which describe the rate of transitioning between states) is too high the likelihood function consequently becomes intractable due to the computational cost of computing the matrix exponential (see Ho et al., 2018; Moler and Van Loan, 2003; Sidje, 1998). Since conventional Bayesian approaches to parameter estimation are no longer feasible, we are motivated to use likelihood-free methods.

### 3.2 Approximate Bayesian Computation

Using a Bayesian framework, the uncertainty about the unknown parameter *θ* with respect to the data *y* can be quantified by sampling from the posterior distribution *π*(*θ|y*) ∝: *π*(*y|θ*)*π*(*θ*); where *π*(*y|θ*) is the likelihood function and *π*(*θ*) is the prior. However, the likelihood function for sufficiently complex models becomes intractable (see examples in biology (Johnston et al., 2016; Vo et al., 2015), ecology (Guillemaud et al., 2010; Toni et al., 2009) and cosmology (Weyant et al., 2013)). Rather than reverting to simpler models with tractable likelihoods, these types of problems can be instead analysed using likelihood-free methods that avoid evaluating the likelihood function.

One popular likelihood-free approach is ABC (Sisson et al., 2018). ABC involves simulating data from the model *x* ~ *f* (·|*θ*) instead of evaluating the intractable likelihood; accepting configurations of *θ* which produce simulated data *x* that is close to the observed data *y*. It can be impractical to compare the full data sets of *x* and *y*, so ABC often relies on reducing the full data sets to summary statistics by some summarising function *S*(·), where the summary statistics for *x* and *y* are denoted *S_x_* = *S*(*x*) and *S_y_* = *S*(*y*), respectively. Provided the summary statistics are highly informative about the model parameters, then *π*(*θ|y*) ≈ *π*(*θ|S_y_*) is a good approximation (Blum et al., 2013). In effect, ABC samples from the approximate posterior:

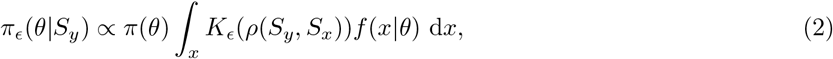

where *ρ*(*S_y_, S_x_*) is the discrepancy function which measures the difference between the two data sets and *K_ϵ_*(·) is the kernel weighting function which weighs *ρ*(*S_y_, S_x_*) conditional on the tolerance *ϵ*. A common choice for the discrepancy function is the Euclidean distance, *ρ*(*S_y_,S_x_*) = ∥*S_y_* — *S_x_*∥_2_, and for the kernel weighting function is the indicator function, 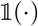, which is equal to one if *ρ*(*S_y_,S_x_*) ≤ *ϵ* and is zero otherwise. The approximate posterior in Equation 2 converges to the posterior conditional on the observed summary (often referred to as the partial posterior) in the limit as *ϵ* → 0 (Beaumont et al., 2002).

To sample from the approximate posterior, commonly ABC-rejection (Pritchard et al., 1999; Tavaré et al., 1997), Markov Chain Monte Carlo ABC (MCMC-ABC) (Marjoram et al., 2003), or Sequential Monte Carlo ABC (SMC-ABC) (Drovandi & Pettitt, 2011; Sisson et al., 2007) algorithms are used. ABC-rejection samples particles from the prior distribution and accepts particles with a discrepancy measure *ρ*(*S_y_,S_x_*) less than the desired tolerance *ϵ*. In cases when the prior distribution is relatively diffuse compared to the posterior density (such as our application), lower acceptance rates are common because particles are predominantly sampled in regions of low posterior density (Sisson et al., 2007). To increase acceptance rates, one could instead use MCMC-ABC which constructs a Markov chain with a stationary distribution identical to the approximate posterior by proposing particles from a carefully-tuned proposal distribution, *θ^i^* ~ *q*(·|*θ*^*i*-1^), and accepting those with probability

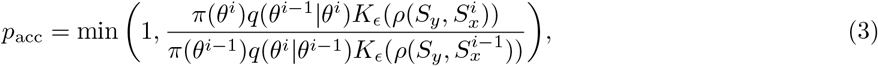

which is based on the Metropolis-Hastings ratio (Hastings, 1970; Metropolis et al., 1953). While MCMC-ABC tends to be more computationally efficient compared to ABC-rejection (Marjoram et al., 2003), it is possible for the Markov chain to spend many iterations in areas of low posterior probability. In our application we found MCMC-ABC to take a considerable effort to tune the proposal distribution while still being computationally cumbersome. However, SMC-ABC or more specifically the SMC-ABC replenishment algorithm (Drovandi & Pettitt, 2011) requires very little tuning comparatively and allows for simulations to be performed in parallel to increase computational efficiency.

The SMC-ABC replenishment algorithm traverses a set of distributions defined by *T* non-increasing tolerance levels *ϵ*_1_ ≤…≤ *ϵ_T_* to sample from the approximate posterior:

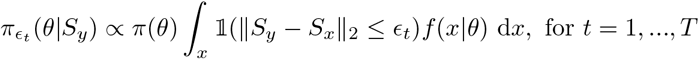

where the first target distribution is constructed by sampling from the prior distribution to attain a collection of parameter values (called particles) and their discrepancies, 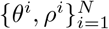. The first tolerance threshold, *ϵ*_1_, is set as the maximum of the set of discrepancies. Thereafter, to propagate particles through the sequence of target distributions, particles are first sorted in ascending order by their discrepancy and the new tolerance is set as *ϵ_t_* = *ρ^N-Nα^* where *N_α_* = ⌊*αN*⌋, *α* is the proportion of particles discarded and ⌊·⌋ is the floor function. Particles, 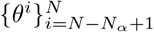, which do not satisfy the new tolerance are discarded and resampled, with replacement, from the remaining particles to replenish the population. To prevent sample degeneracy (too many duplicated particles), resampled particles are then perturbed according to an MCMC kernel *R_t_* times with an invariant distribution given by the current approximate posterior *π_ϵt_* (*θ*|*S_y_*). In each of the *R_t_* iterations, the proposed particles are drawn from an automatically tuned proposal distribution *θ^i^* ~ *q*(·|*θ*^*i*-1^) and accepted with probability 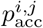 (Equation 3) where *i* denotes the *i*th particle and *j* the *j*th MCMC iteration. To ensure sample diversity, *R_t_* can be dynamically set based on the overall MCMC acceptance rate, 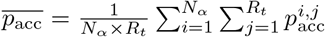, such that there is a 1 — *c* chance that all particles are moved at least once and is given by

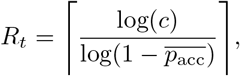

where the ceiling function ⌈·⌉ is used to be conservative and an estimate for 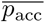 is calculated from *R*_*t*-1_/2 pilot MCMC iterations. A popular choice for the proposal distribution is the multivariate normal distribution, 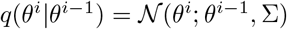, where Σ is the tuning parameter which can be adaptively tuned from the 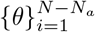 particles which are already distributed according to the current target distribution. The algorithm finally stops once the overall MCMC acceptance rate, 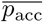, is unreasonably low (≤ 1%) or the desired tolerance threshold is reached. For the two tuning parameters, Drovandi and Pettitt (2011) suggest setting *α* = 0.5 and *c* = 0.01. The SMC-ABC replenishment algorithm is presented in Algorithm 1. A crucial limitation of ABC methods is the curse of dimensionality, where despite the addition of more data, the approximation to the posterior can become distorted as a result of the discrepancy between observed data and simulated data *ρ*(*S_y_,S_x_*) naturally increasing with the dimension (Beaumont et al., 2002). In applications where increasing the dimension of the summary statistics cannot be avoided, the discrepancy between observed and summary statistics can be accounted for, at least approximately, with regression adjustment (Beaumont et al., 2002; Blum et al., 2013). Regression adjustment involves explicitly modelling the parameters against the discrepancy between observed and simulated data. Assume for the moment that *θ* is a scalar parameter. Consider the following regression model

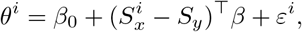

where *i* = 1,…,*N* is the parameter sample index, *β* is the regression coefficients, *β*_0_ is the intercept and *ε^i^* is the error term. Estimates for *β*_0_ and *β* can be computed by minimising the weighted least squares criterion 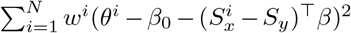. Here we choose to use the popular Epanechnikov weighting function (Epanechnikov, 1969), defined as 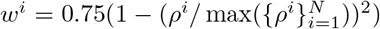, but other weighting functions could also be used. Using the estimated regression coefficients 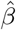, we then make the adjustment

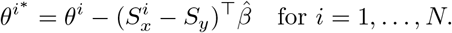

The adjusted sample 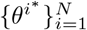 can often give a more accurate approximation of the posterior. To ensure that the adjusted parameters remain within the support of the prior distribution (if bounded), Hamilton et al. (2005) suggest transforming parameter values before applying the regression adjustment. We use a logit transformation, 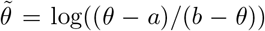, where *a* and *b* are the respective lower and upper bounds of the prior. Given that we have a vector of parameters, we apply a regression adjustment to each component of the parameter vector separately.

**Algorithm 1.**
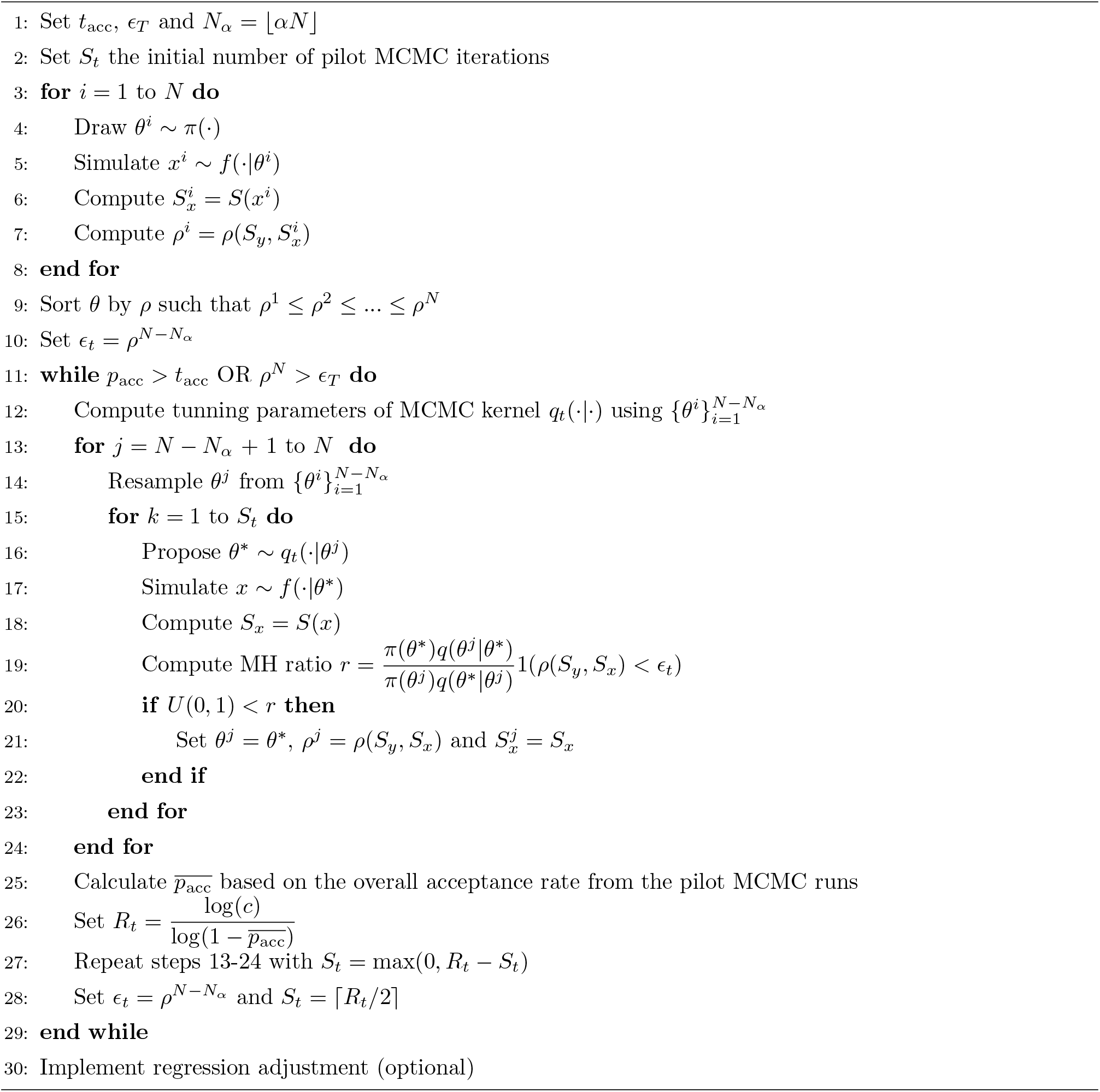
SMC-ABC (Drovandi & Pettitt, 2011)

### 3.3 Prior Knowledge

To quantify the cell cycle transition rates *R* ∈ {*R_r_, R_y_, R_g_*}, commonly the cell doubling time *t_d_* is used, where *t_d_* = ln(2)/*R*. Estimates for the cell doubling time for melanoma cells range from 16-46 h (Haass et al., 2014; Simpson et al., 2020). Furthermore, Simpson et al. (2020) estimate the average time 1205Lu FUCCI-transduced melanoma cells spend in the G1 and S/G2/M phases to be between 8-30 h and 8-17 h, respectively, and estimate the transition rates between phases to be in the range from 0.03-0.125/h. Therefore, we propose that our prior information of the transition rates is uniform over the range 0 — 1/h to be conservative.

Cell diffusivity, *D*, the measurement of motility rate for particles undergoing random diffusive migration, can be used to quantify the cell motility rate *M* ∈ {*M_r_, M_y_, M_g_*}, by *D* = *M*Δ^2^/4 (Codling et al., 2008), where Δ is the cell diameter. Empirical evidence finds estimates for cell diffusivity to range from 0 — 3304 *μ*m^2^/h (Cai et al., 2007; Maini et al., 2004; Treloar et al., 2013). Furthermore, Simpson et al. (2018) suggest the cell diffusivity to be approximately 400 *μ*m^2^/h, so that the rates are approximately 4/h. We propose that the prior information of the motility rates to be uniform over the range 0 — 10/h; attributing the larger interval to the greater variation of cell diffusivity estimates in existing literature.

## 4 Results

For SMC-ABC we generate samples from the approximate posteriors using *N* = 1000 particles. From preliminary trials, we found it more useful to use the overall MCMC acceptance rate as the stopping rule for the SMC-ABC algorithm and adopt the sensible choice for the target acceptance rate as *t_acc_* = 1% and *ϵ_max_* = 0 for the target tolerance.

### 4.1 Developing summary statistics and validation with synthetic data

The accuracy and precision of ABC methods in approximating the posterior distribution is sensitive to the quality of the summary statistics used (Beaumont et al., 2002). We first trial and validate different summary statistics with multiple synthetic data sets such that the true parameter values are known. In this way, we are able to compare the performance of different summary statistics and determine which are the most effective. While trying to replicate the environment of the experimental data as close as possible, such as domain size, boundary conditions and initial number of cells, we do not calibrate the initial location of cells but rather randomly distribute the cells within a 200 *μ*m by 1745.35 *μ*m region on either side of the scratch. We attain the initial cell counts of red, yellow, and green cells by using the procedure outlined in section 4.2 (steps 1-4) and report them here to be 119, 35 and 121, respectively.

In our analysis of the simulation model, agents with relatively higher transition rates were found to correspond to lower population sizes, and vice versa. Therefore, we use the number of agents in each population (*N_r_, N_y_, N_g_*) at the end of the experiment as summary statistics for the transition rates *R_r_, R_y_, R_g_*, respectively. We test the suitability of this summary statistic on four synthetic data sets produced by varying the transition rates amongst biologically plausible values and keeping the motility rates known and constant. The four parameter configurations we choose to generate the synthetic data sets are *θ* = {(0.04, 0.17, 0.08, 4, 4, 4), (0.25, 0.15, 0.22, 4, 4, 4), (0.12, 0.07, 0.03, 4, 4, 4), (0.3, 0.36, 0.28, 4, 4, 4)}. In Appendix B we present the marginal posterior distributions produced and confirm the suitability of this summary statistic.

For the motility parameters we explore and compare two sets of summary statistics, namely cell density and cell trajectory data. Of these two data sets, cell density data is desirable due to less manual effort needed to generate the data while cell trajectory data could offer more information but is more challenging to collect. For the cell density data, we first segment the imaged region at the end of the experiment (*t* = 48 h) directly down the centre of the image in the *y* direction and calculate the median position and interquartile ranges of the red, yellow, and green agent populations in the *x* direction for cells on the left and right sides. For the cell trajectory data, we record the distance the cell travels through each cell phase until the cell returns to the initial phase or the simulation is terminated. We select cells to be tracked provided that the cell is initially in G1 (red) phase and the cell is located on the leading edge of the cell monolayer toward the gap in the scratch assay. In Appendix B, we draw samples from the posterior distribution using 10, 20, 30, 40 and 50 cell trajectories with four synthetic data sets generated from *θ* = {(0.04, 0.17, 0.08, 4, 4, 4), (0.04, 0.17, 0.08, 2, 5, 8), (0.04, 0.17, 0.08, 8, 2, 5), (0.04, 0.17, 0.08, 5, 8, 2)}. Our analysis concludes 20 cells to be the minimum number of cells necessary to produce a well defined posterior. Using the same four synthetic data sets we also attempt to draw samples from the posterior distribution using cell density data in Appendix B, however, we found the motility parameters to be non-identifiable when the transition rates are held constant.

We now combine the summary statistics formulated to estimate the cell cycle transition and motility rates together with four synthetic data sets. Due to the consistency in estimates for cell cycle transition rates in existing literature (see Haass et al., 2014; Simpson et al., 2020), we adopt estimates for the cell cycle transition rates from Haass et al. (2014) for all four parameter configurations. Since estimates for motility rates have been reported to vary by two orders of magnitude (see Cai et al., 2007; Maini et al., 2004; Treloar et al., 2013), we choose to vary the motility rates over the range of the prior for the four parameter configurations. That is, we generate four synthetic data sets with *θ* = {(0.04, 0.17, 0.08, 4, 4, 4), (0.04, 0.17, 0.08, 2, 5, 8), (0.04, 0.17, 0.08, 8, 2, 5), (0.04, 0.17, 0.08, 5, 8, 2)}. In Figure 3 we present the marginal posterior distributions produced when using the number of cells in each subpopulation and cell density data as summary statistics. In Figure 4 we present the marginal posterior distributions when using cell trajectory data in place of cell density data. Again, we see that the motility estimates are unidentifiable when cell density data is used while both cell cycle transition and motility estimates are identifiable when cell trajectory data is included. Furthermore, it is clear from the concentration of the marginal posterior distributions around the true parameter values (dashed line) in Figure 4 that the cell count and cell trajectory data are highly informative about the transition and motility parameters, respectively. We note that the precision of these distributions is greater for the cell cycle transition parameters than the motility parameters. Importantly, these results show for the first time that practical parameter inference on both transition and motility parameters of a FUCCI scratch assay experiment using Bayesian inference techniques is possible. These results justify the choice of the Markov process model compared to simpler continuum models which do not give insight into cell trajectory data (see Simpson et al., 2020).

**Figure 3:**
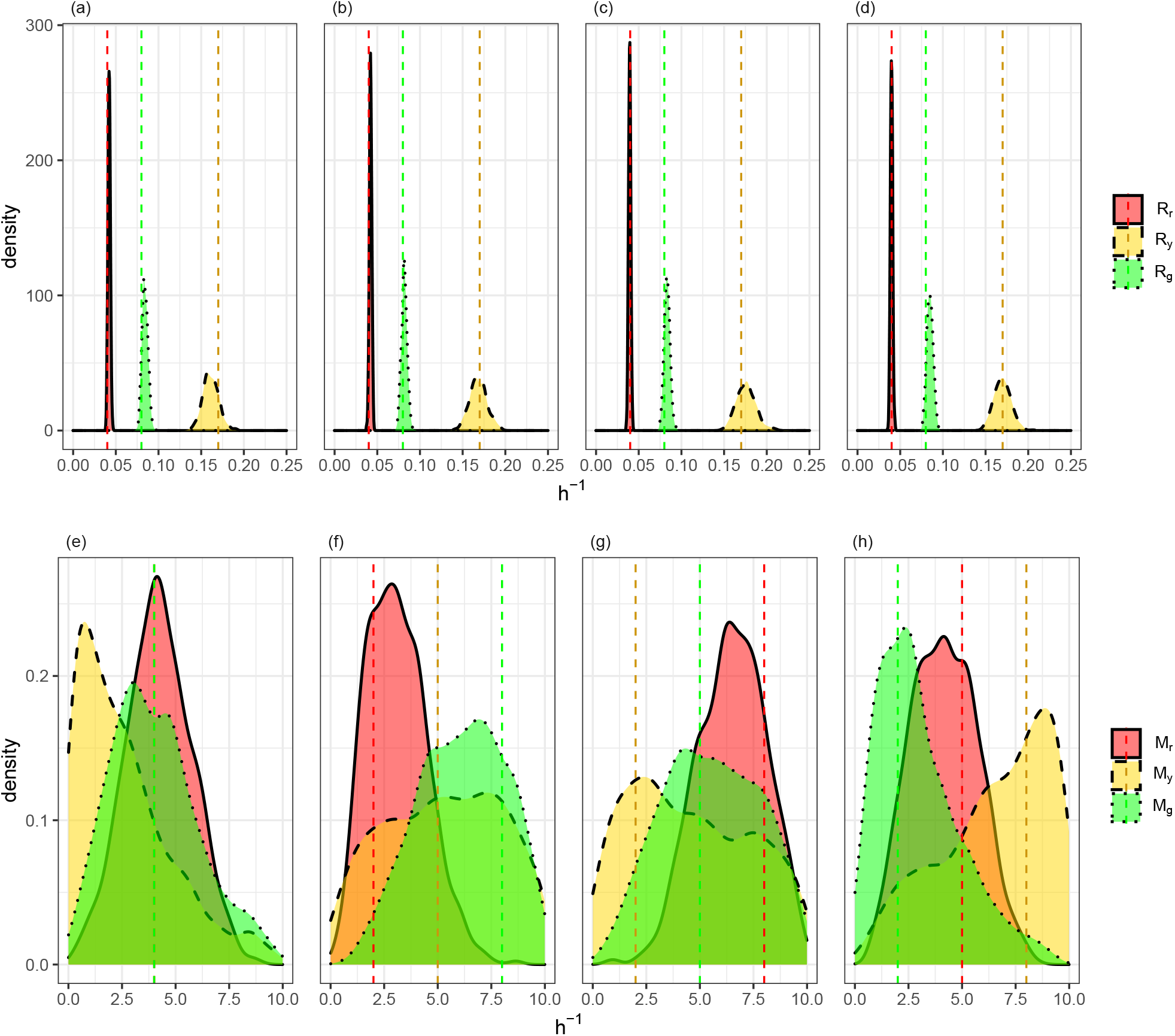
Estimating cell cycle transition and cell motility parameters, *θ* = (*R_r_, R_y_, R_g_, M_r_, M_y_, M_g_*), with the number of cells in each phase at *t* = 48 h and cell density data as summary statistics across several synthetic data sets. Synthetic data sets were produced from simulations with true parameter values indicated by dashed line. (a,e) Estimated marginal posteriors produced with *θ* = (0.04, 0.17, 0.08, 4, 4, 4). (b,f) Estimated marginal posteriors produced with *θ* = (0.04, 0.17, 0.08, 2, 5, 8). (c,g) Estimated marginal posteriors produced with *θ* = (0.04, 0.17, 0.08, 8, 2, 5). (d,h) Estimated marginal posteriors produced with *θ* = (0.04, 0.17, 0.08, 5, 8, 2).

**Figure 4:**
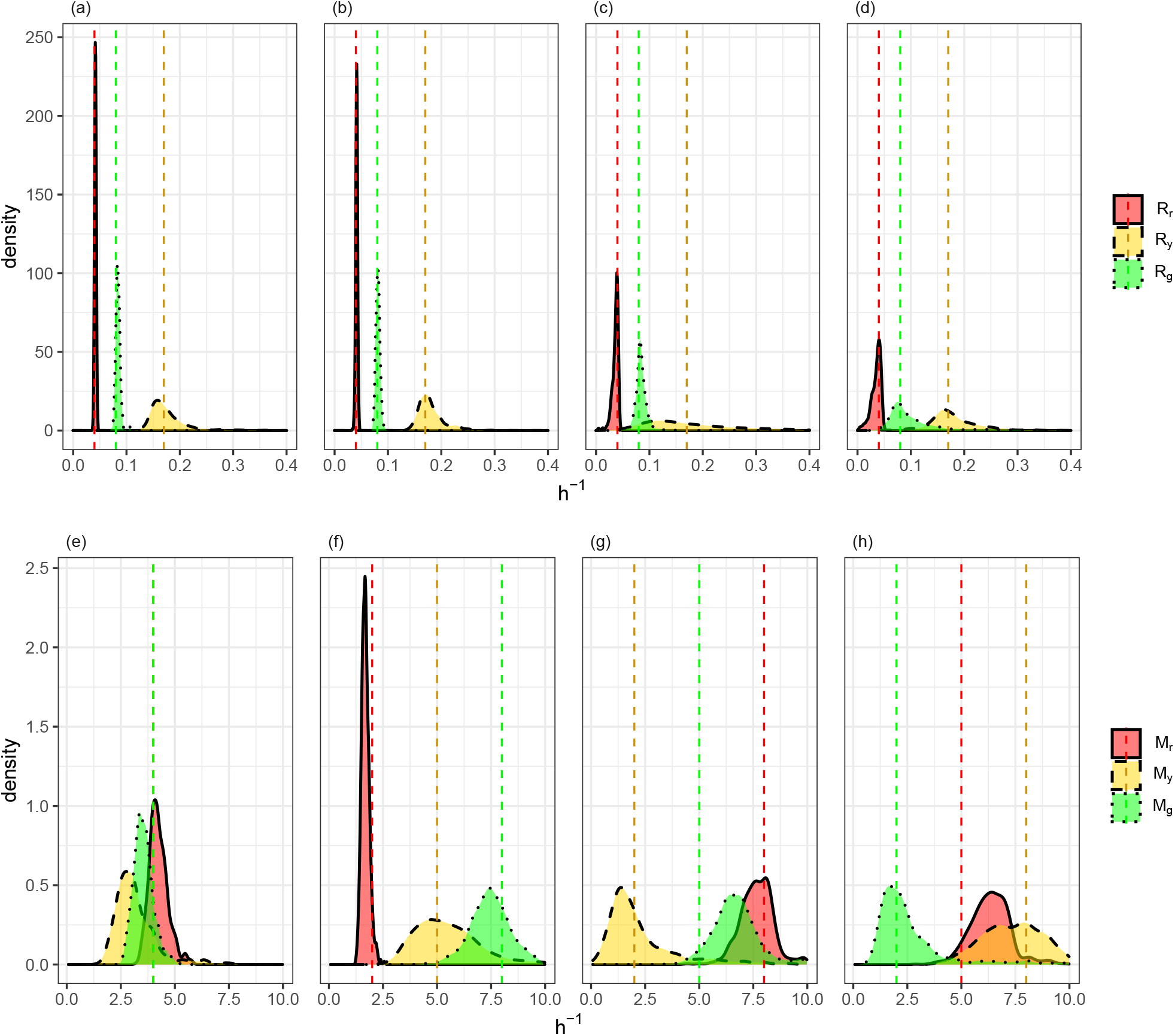
Estimating cell cycle transition and cell motility parameters, *θ* = (*R_r_, R_y_, R_g_, M_r_, M_y_, M_g_*), with the number of cells in each phase at *t* = 48 h and cell tracking data as summary statistics across several synthetic data sets. Synthetic data sets were produced from simulations with true parameter values indicated by dashed line. (a,e) Estimated marginal posteriors produced with *θ* = (0.04, 0.17, 0.08, 4, 4, 4). (b,f) Estimated marginal posteriors produced with *θ* = (0.04, 0.17, 0.08, 2, 5, 8). (c,g) Estimated marginal posteriors produced with *θ* = (0.04, 0.17, 0.08, 8, 2, 5). (d,h) Estimated marginal posteriors produced with *θ* = (0.04, 0.17, 0.08, 5, 8, 2).

### 4.2 Image analysis of experimental data

We analyse the experimental images using *ImageJ* (Rueden et al., 2017) to record the Cartesian coordinates of cells. Of primary interest is processing the initial frame such that we can replicate the experimental settings as accurately as possible in the simulation but we also repeat this procedure for the final frame to retrieve the final cell counts and cell density data, which we use as summary statistics. The process is as follows:

1. Step 1: *Read in image:* File > Open > *select image* (Figure 5 (a)).
2. Step 2: *Convert image to 8-bit:* Image > Type > 8-bit (Figure 5 (b)).
3. Step 3: *Identify cell edges:* Convert the image to black and white (Process > Binary > Convert to Mask) and then distinguish conjoined cells (Process > Binary > Watershed) (Figure 5 (c)).
4. Step 4: *Compute Cartesian coordinates:* Analyze > Analyze Particles…> OK.

**Figure 5:**
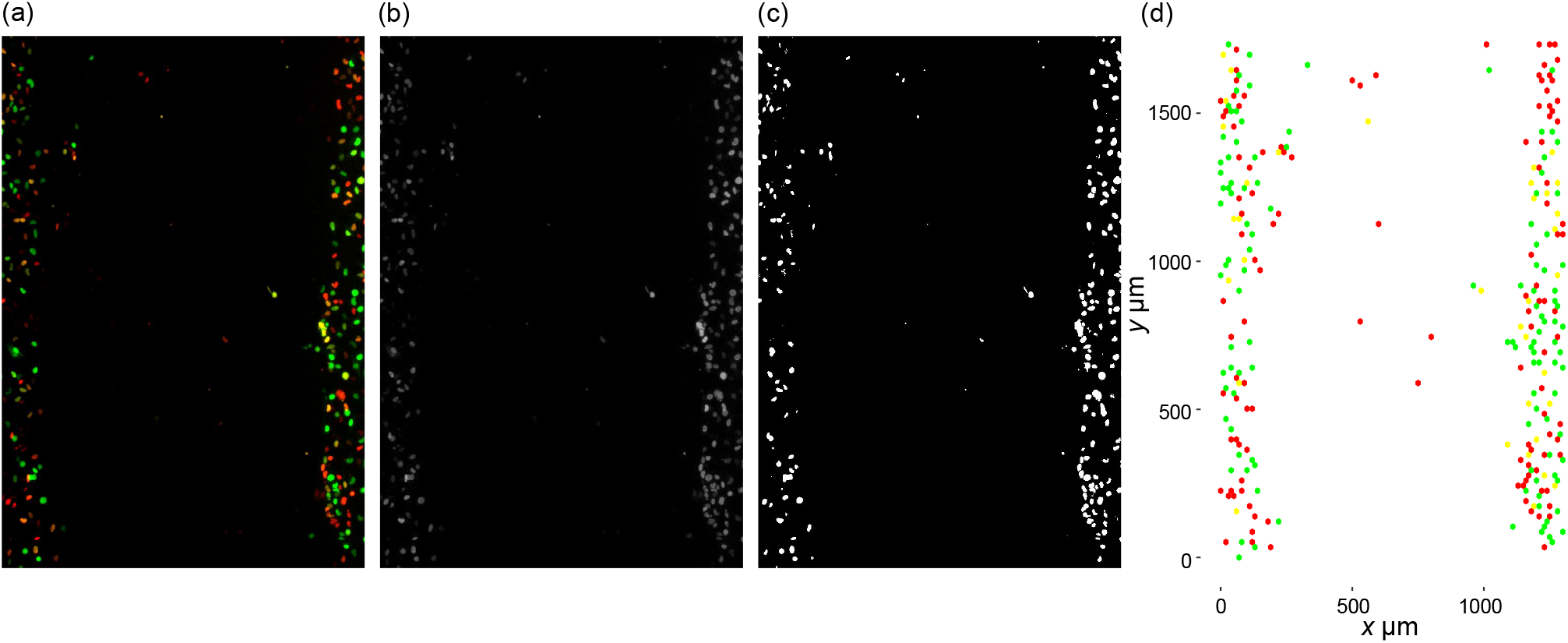
*ImageJ* procedure. (a) Original image loaded (WM983C FUCCI-transduced melanoma cells). (b) Image after compression to 8-bit. (c) Image after converting to black and white and watershedding. (d) Simulation initial geometry recovered from data processing of WM983C FUCCI-transduced melanoma cells in *ImageJ* and *R*

A limitation of using the watershed tool is that we must convert the image to black and white. In doing so, we lose the cell phase identity associated with the cell coordinates recovered from *ImageJ*. To overcome this, we use *R* (R Core Team, 2020) to retrieve the RGB decimal color code and Cartesian coordinates of pixels. Matching pixel coordinates recovered from *R* and the coordinates of the centroid of the cells recovered in *ImageJ*, we create a data set of cell coordinates and their associated RGB decimal codes. To classify the RGB coordinates into one of the three cell cycle phases we use the conditions outlined in Table 1.

**Table 1:**
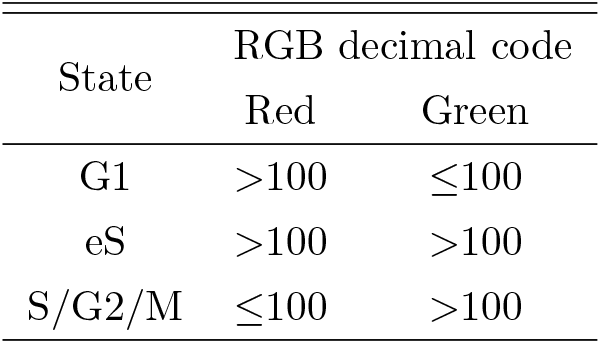
Cell phase classification rule using RGB decimal codes.

To extract summary statistics from the experimental data, we repeat the image processing procedure previously outlined above with the final frame (*t* = 48 h) and extract the final cell counts and cell density data. Additionally, we extract cell trajectory data by processing the entire sequence of still images in *ImageJ* with the “Multi-point” tool to manually track cell coordinates between frames. We use a similar process as before to identify cell phases in these summary statistics using *R* and present them in Table 2 and the cell trajectory data in Figure 6.

**Figure 6:**
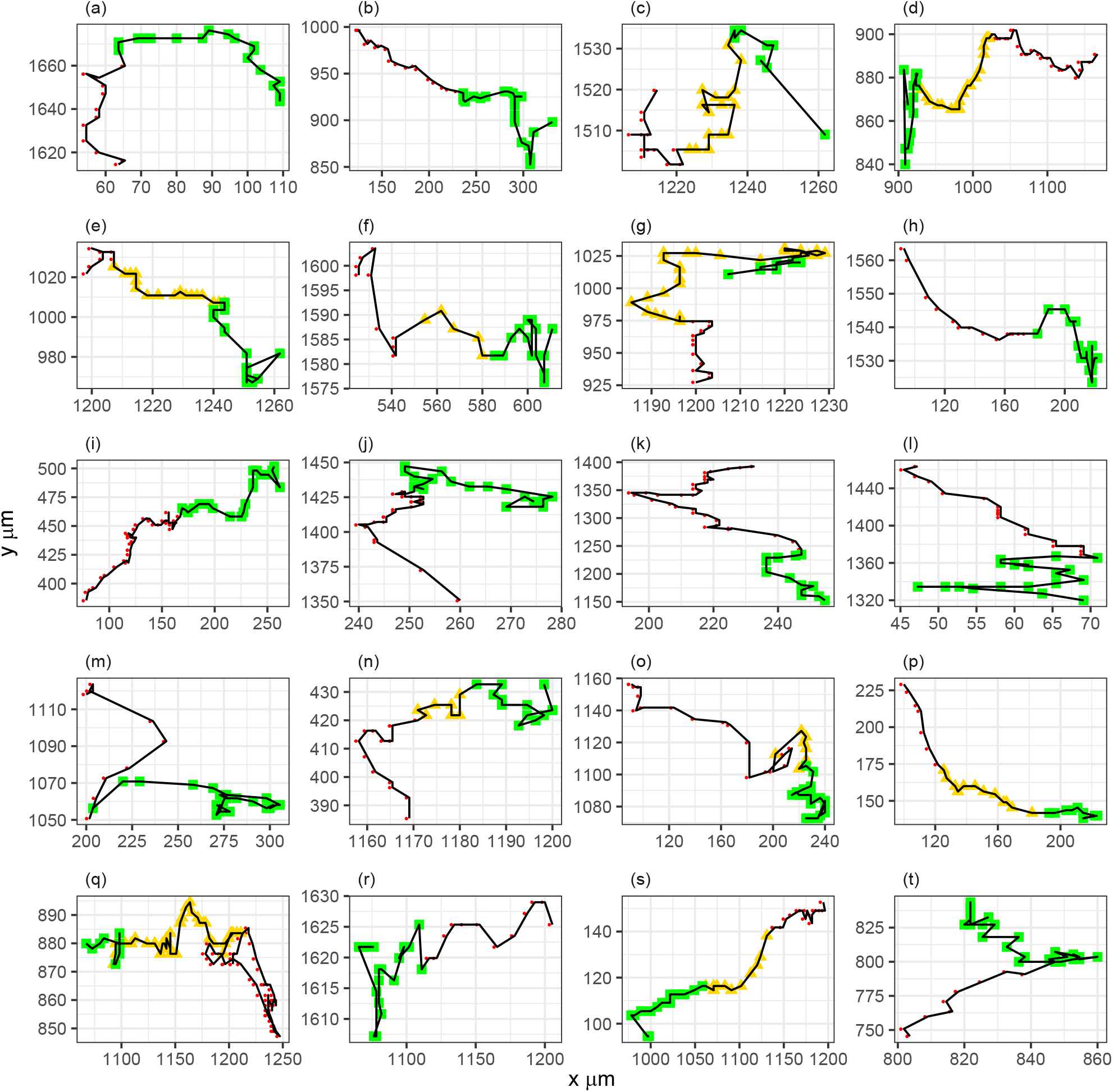
Trajectories of WM983C FUCCI-transduced melanoma cells where each box ((a)-(t)) corresponds to one of the twenty cell trajectories. Tracking begins in red phase (red circles) then progresses through the yellow phase (yellow triangles) and is terminated at end of green phase (green squares).

**Table 2:**
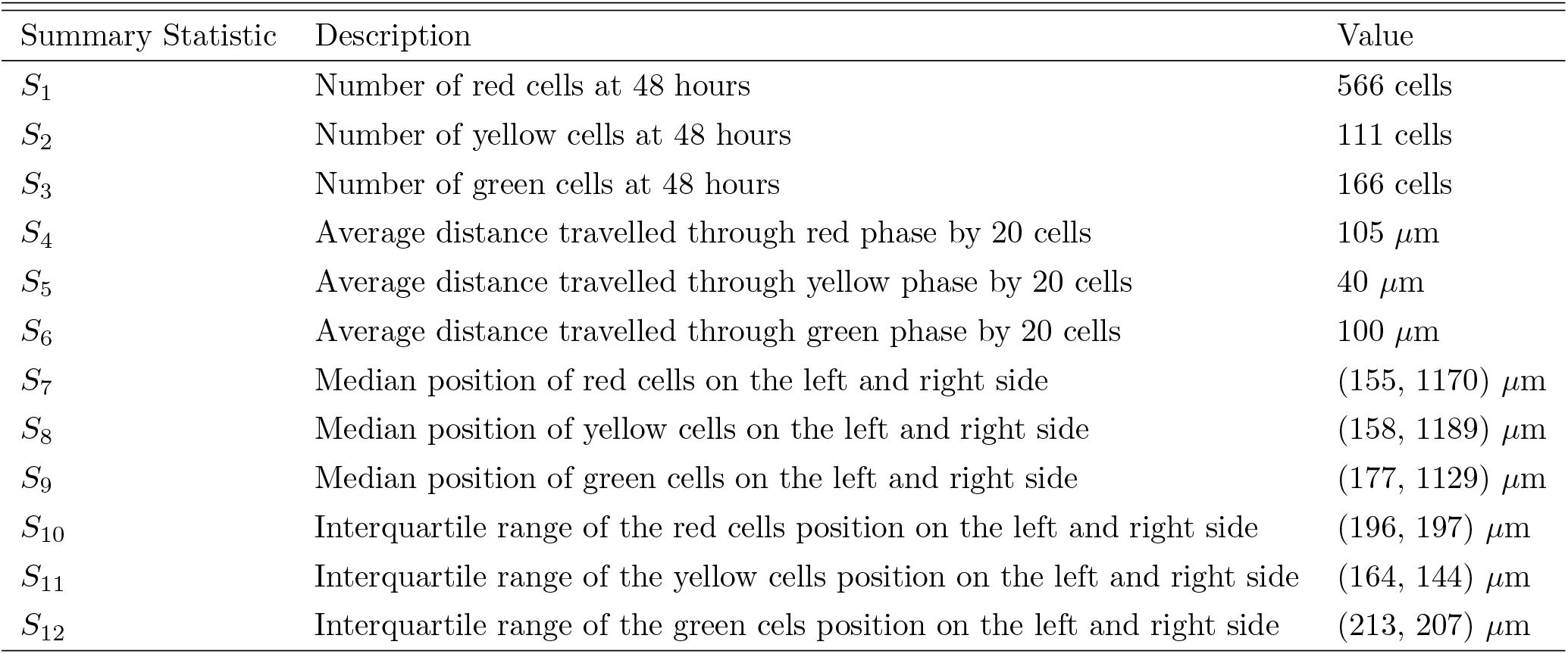
Observed summary statistics of WM983C FUCCI-transduced melanoma cells

Finally, we calibrate the hexagonal lattice used in the simulation model with the data set of Cartesian coordinates recovered previously by rearranging Equation 1 to find their associated lattice row and column indices denoted

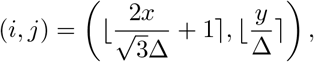

where ⌊·⌉ rounds to the nearest integer. We treat the rare instances (<1%) where multiple coordinates are mapped to the same lattice space as duplicated values and omit them rather than place them on the next closest lattice site. The result from this translation of data is presented in Figure 5 (d). We repeat this process for the initial frame of the cell trajectory data to identify the starting position. However, due to manually tracking cell trajectories, often the coordinate retrieved was not centred on the cell which in some cases caused the starting position to be mapped to an unoccupied lattice site. We intervene prior to transforming the starting position and adjust the coordinate values to the closest occupied lattice site which is chosen such that the radial distance between the coordinate and the lattice site is minimised.

### 4.3 Estimating Model Parameters with Experimental Data

After calibrating the simulation to the experimental data of WM983C FUCCI-transduced melanoma cells, we first attempt to sample from the posterior distribution using the number of cells in each subpopulation and cell density data (summary statistics *S*_1_ to *S*_3_ and *S*_7_ to *S*_12_ in Table 2, respectively) and present the samples from the posterior distribution in Figure 7. Consistent with results found in Section 4.1 and Simpson et al. (2020), estimates for the motility rates are non-identifiable when cell density data is used.

**Figure 7:**
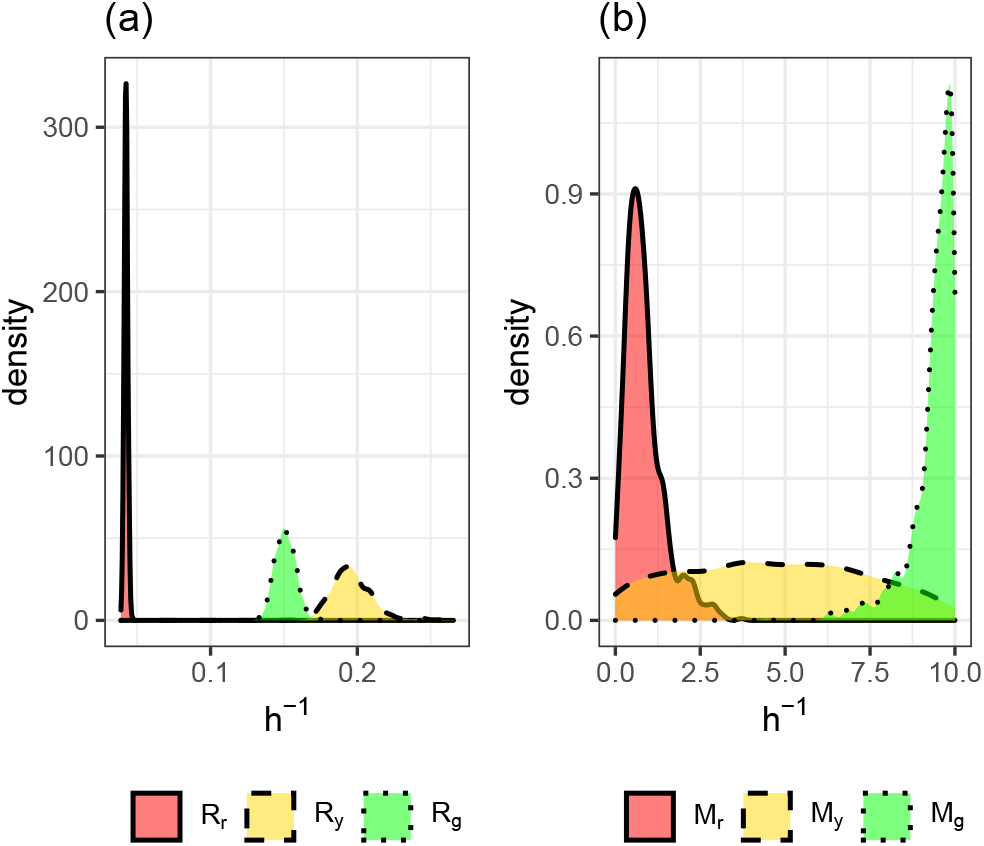
Marginal posterior distributions using number of cells in each subpopulation and cell density data. (a) Marginal posterior distributions for transition rates of WM983C FUCCI-transduced melanoma cells. (b) Marginal posterior distributions for motility rates of WM983C FUCCI-transduced melanoma cells.

Next, we attempt to sample from the posterior distribution using the number of cells in each subpopulation and cell trajectory data (summary statistics *S*_1_ to *S*_3_ and *S*_4_ to *S*_6_ in Table 2, respectively). We present the marginal posterior distributions produced in Figures 8 (a)-(b) along with the mean, standard deviation, (2.5%, 50%, 97.5%) quantiles, and the coefficient of variation (CV) in Table 3. The identifiability in the transition and motility parameters clearly shows the benefits of using cell tracking data as the distributions are unimodal and concentrated. We estimate the cell cycle transition rates to be between 0.0411 — 0.193/h which is consistent with estimates in existing literature (Haass et al., 2014; Simpson et al., 2020). Our estimates for cell motility were found to range between 0.316 — 1.12/h which corresponds to estimates of cell diffusivity between 31.6 and 112 *μ*m^2^/h which is reasonable considering the degree of uncertainty in existing estimates which can vary between 0 and 3304 *μ*m^2^/h (Cai et al., 2007; Maini et al., 2004; Treloar et al., 2013). The precision in parameter estimates can be quantified by the CV which is a standard measure for the dispersion of data around the mean. Using the CV, the dispersion in the transition rates range from 2.65 — 5.31% and the motility rates range from 10.9 — 18.4%. To validate the parameter estimates recovered, we also present the posterior predictive distributions for the summary statistics retained from each parameter value in the posterior in Figures 8 (c)-(d). These distributions are formed by plotting the distribution of simulated summary statistics produced from the posterior samples and is compared to the observed summary statistics (dashed line). These results suggest that the Markov process model developed by Simpson et al. (2018) is appropriate as it is able to recover the observed summary statistics of the experimental data.

**Figure 8:**
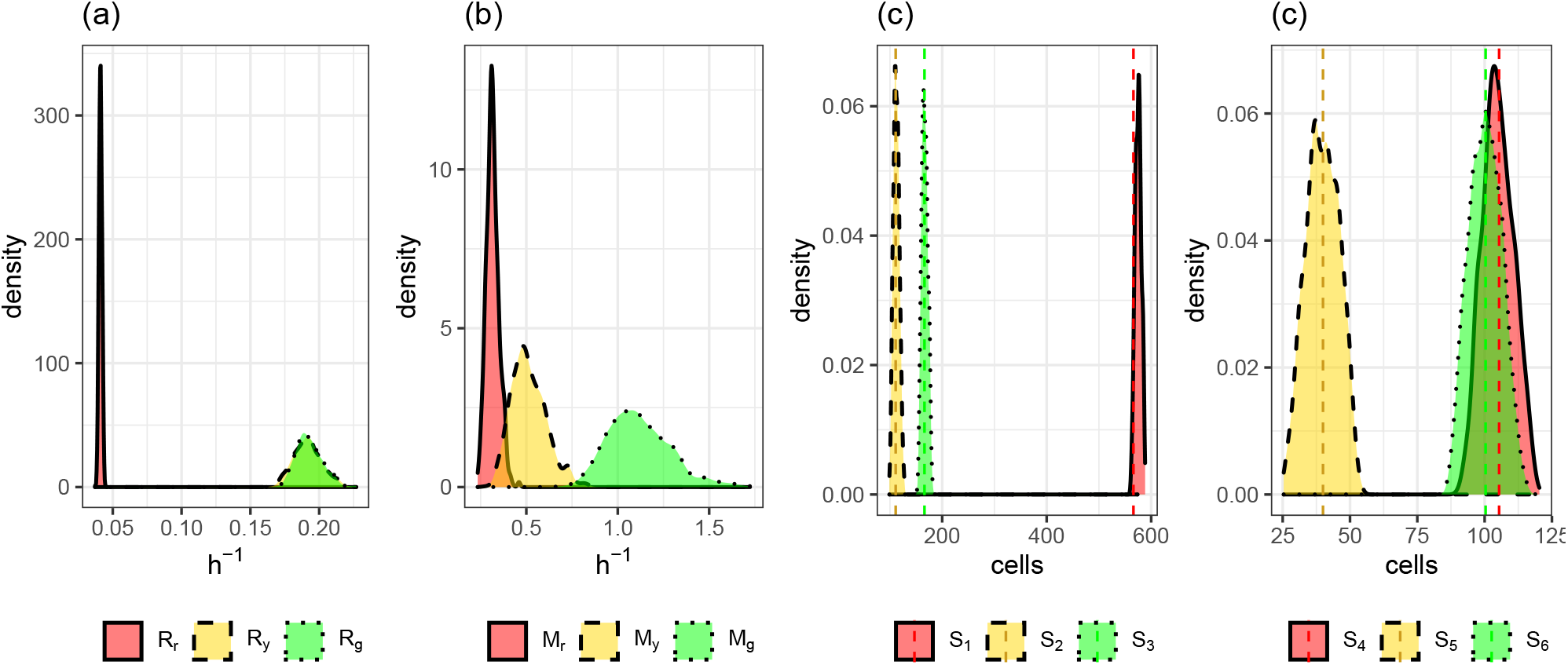
Marginal posterior distributions using number of cells in each subpopulation and cell trajectory data. (a) Marginal posterior distributions for transition rates of WM983C FUCCI-transduced melanoma cells. (b) Marginal posterior distributions for motility rates of WM983C FUCCI-transduced melanoma cells. (c) Distribution of simulated summary statistics (informative of transition rates) compared to observed summary statistics (dashed line). (d) Distribution of simulated summary statistics (informative of motility rates) compared to observed summary statistics (dashed line).

**Table 3:** Posterior summaries (3 significant figures): mean, standard deviation, (2.5%, 50%, 97.5%) quantiles, and the coefficient of variance (CV).

| Parameter | Mean | Std. Dev. | (2.5%, 50%, 97.5%) | CV (%) |
| --- | --- | --- | --- | --- |
| $R_r$ | 0.0411 | 0.00109 | (0.039, 0.0411, 0.0432) | 2.65 |
| $R_y$ | 0.192 | 0.0102 | (0.173, 0.191, 0.214) | 5.31 |
| $R_g$ | 0.193 | 0.00957 | (0.177, 0.192, 0.213) | 4.96 |
| $M_r$ | 0.316 | 0.0343 | (0.256, 0.313, 0.385) | 10.9 |
| $M_y$ | 0.514 | 0.0945 | (0.353, 0.502, 0.725) | 18.4 |
| $M_g$ | 1.12 | 0.169 | (0.836, 1.10, 1.48) | 15.1 |

## 5 Discussion

In this study, we calibrate the 2D hexagonal-lattice random walk model developed by Simpson et al. (2018) to scratch assay data where the cell cycle is revealed in real time using FUCCI technology. While this model is well suited to describing the stochastic nature of cell proliferation and migration, the likelihood function consequently becomes intractable. This makes conventional Bayesian approaches to parameter inference infeasible. We resort to using the class of Bayesian methods known as ABC which bypass evaluating the likelihood function. After evaluating the appropriateness of different ABC algorithms we find the SMC-ABC replenishment algorithm developed by Drovandi and Pettitt (2011) to be suitable.

The accuracy of ABC methods in approximating the posterior distribution is sensitive to the quality of the summary statistics used (Beaumont et al., 2002). We trial various summary statistics with multiple synthetic data sets to determine which summary statistics are the most informative. We find using the number of cells in each cell cycle phase at the end of the experiment to be highly informative about the cell cycle transition rates. We trial and compare two sets of summary statistics for the motility parameters: the median position and interquartile range of the cells in the x direction on the left and right side of the scratch assay (which we refer to as cell density data); and the average distance travelled through each cell phase by 20 individual cells (which we refer to as cell trajectory data). Using these two sets of summary statistics in conjunction with the cell cycle transition summary statistics, we attempt to draw samples from the posterior distribution using the SMC-ABC replenishment algorithm with multiple biologically plausible synthetic data sets. We find that when using cell trajectory data as summary statistics the parameter estimates were identifiable; however this was not the case when cell density was used. Importantly, this is the first time practical parameter identifiability for both cell cycle transition and motility has been successfully conducted with fluorescent cell cycle labelling scratch assay experiments.

We extend on the work of previous studies (Simpson et al., 2020; Simpson et al., 2018) by calibrating our model to real data and performing Bayesian inference. Using experimental data of WM983C FUCCI-transduced melanoma cells, we estimate the approximate posterior using the SMC-ABC algorithm with our cell cycle transition rate summary statistics and our two sets of motility summary statistics. Under the experimental setting, our results again find the estimates for the motility parameters to be non-identifiable when cell density data is used but identifiable when cell trajectory data is used. These results are consistent with Simpson et al. (2020) and justify the motivation to use a more complex model capable of generating multiple data types. When using the number of cells in each subpopulation and cell trajectory data, we find estimates for the average cell cycle transition rates to range between 0.0411 — 0.193/h and estimates for average cell motility to range between 0.316 — 1.12/h. We quantify the precision of these estimates through the CV which is a standard measure of dispersion about the mean. We find the CV to be suitably small for all parameters as it ranges from 2.65 — 5.31% and 10.9 — 18.4% for the transition and motility marginal posteriors, respectively. To validate our results we also draw samples from the posterior predictive to determine whether the simulated data sets recovered accurately reflect the observed data sets. These results confirm that the model and summary statistics are recovering the underlying mechanisms present in the experiment.

Now that the recovery of precise parameter estimates from a fluorescent cell cycle labelling model has been demonstrated, further models can be built which are more biologically realistic. For instance, the Markov process model we used in this study describes a discrete exclusion based random walk on a 2D hexagonal lattice. However, a more biologically realistic and meaningful model would incorporate a three-dimensional (3D) environment (e.g. Jin et al., 2021). By constraining our model to a 2D hexagonal lattice, we ultimately omit realistically modelling: the spatial supply of oxygen, nutrients and drugs; the orientation in 3D space; and interactions with the extracellular matrix (Beaumont et al., 2014; Smalley et al., 2006). Although, increasing model complexity tends to require additional parameters in the model which in some applications may render ABC methods ill suited to inference due to their poorer performance in higher dimensions (Fearnhead & Prangle, 2012). Such modeling and inference implications would need to be considered in future work. Nevertheless, we demonstrate that the 2D stochastic model developed by Simpson et al. (2018) is able to recover key features of the experimental data set we examined and can be used to provide a quick and inexpensive alternative to *in vitro* experiments.

## Acknowledgments

We would like to acknowledge the services of Queensland University of Technologies High Performance Computing (QUT HPC) for allowing the code used within this study to be ran on their servers. Additionally, MJC would like to thank funding provided by MJS and CD.

## Data accessibility

Rcpp and MATLAB implementations of the simulation model and SMC-ABC algorithm along with the data used in this study are available at https://github.com/michaelcarr-stats/FUCCI.

## Authors’ contributions

All authors designed the research. MJC performed the research, wrote the manuscript, produced the figures, and implemented the computational algorithms used in this study. All Authors edited and approved this manuscript.

## Appendix

### A Gillespie Algorithm

**Algorithm 2.**
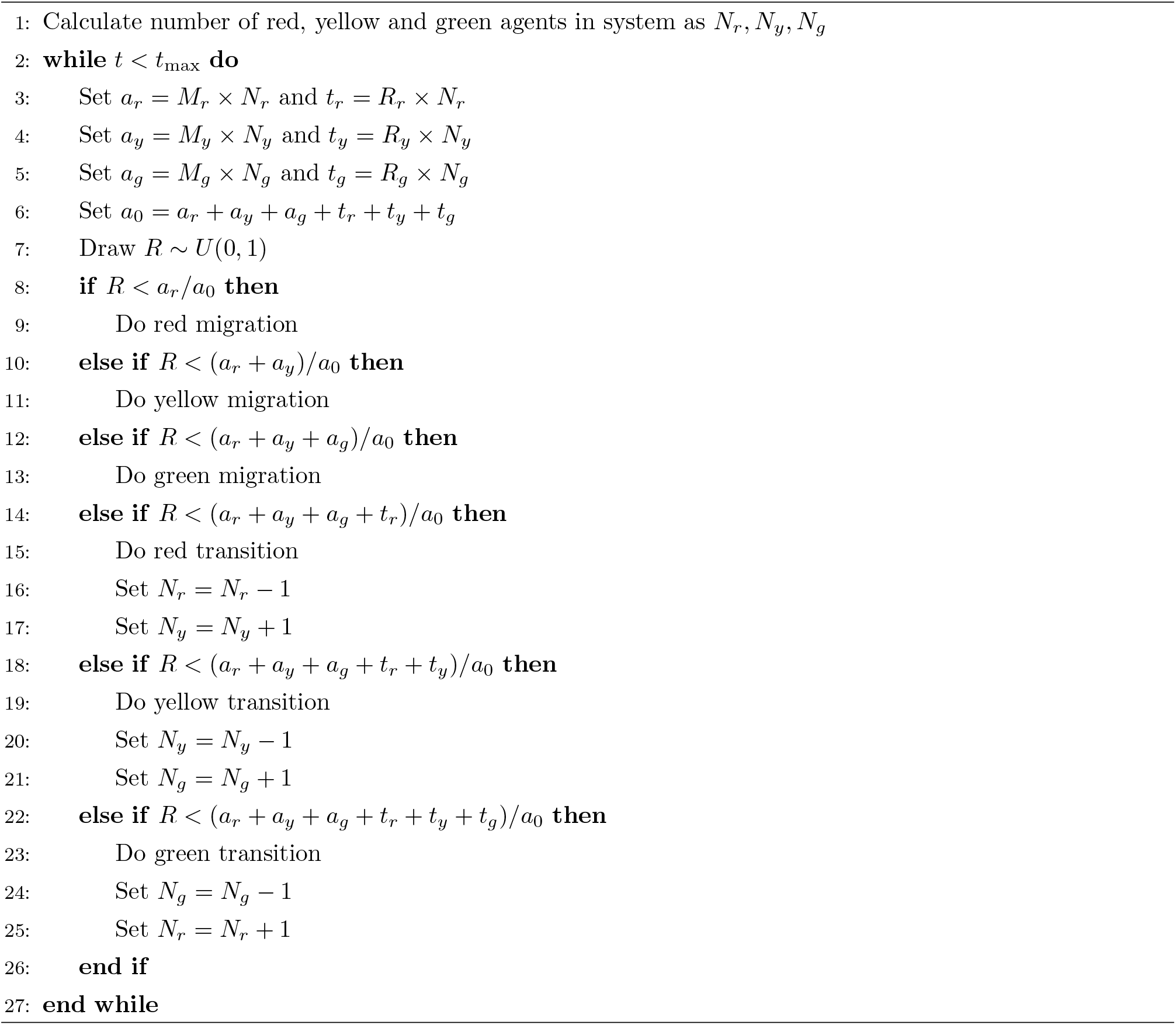
Simulation Model utilising Gillespie Algorithm (Gillespie, 1977) with input parameter *θ* = (*R_r_, R_y_, R_g_, M_r_, M_y_, M_g_*)

### B Developing Summary Statistics

#### Cell Cycle Transition Rates

The summary statistic we explore for the cell cycle transition rates is the number of agents within each phase of the cell cycle (red, yellow, green) at time t = 48 hours. We believe that the count of each cell type to be a good choice because the transition rates will only influence the number of cells. Therefore, we expect higher relative transition rates to correlate to lower cell counts and vice versa. For simplicity, we will assume the motility rates to be known and equal while we estimate the cell cycle transition parameters using multiple synthetic data sets generated from *θ* = {(0.04, 0.17, 0.08, 4, 4, 4), (0.25, 0.15, 0.22, 4, 4, 4), (0.12, 0.07, 0.03, 4, 4, 4), (0.3, 0.36, 0.28, 4, 4, 4)}. Using the SMC-ABC algorithm with the same summary statistics for the simulated data, we present the marginal posterior distribution of the cell cycle transition rates in Figure 9. Since the posterior distributions are all centred on the “true” value we confirm the summary statistics suitability at identifying the cell cycle transition parameters when the motility parameters are held constant and known.

**Figure 9:**
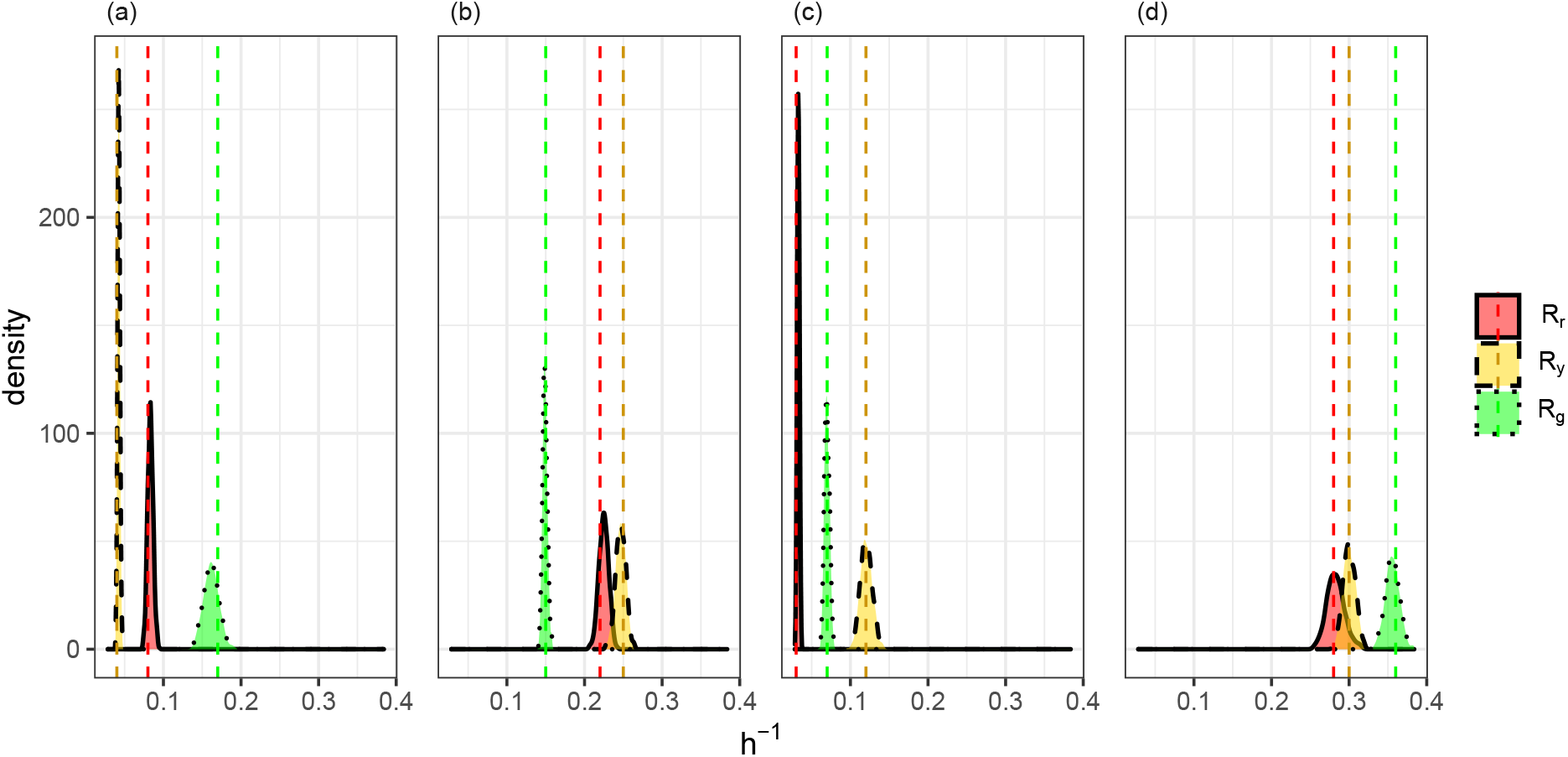
Using number of cells in each cell cycle phase as summary statistics for the transition rates. (a)-(d) Posterior distributions produced using synthetic data sets generated from *θ* = {(0.04, 0.17, 0.08, 4, 4, 4), (0.25, 0.15, 0.22, 4, 4, 4), (0.12, 0.07, 0.03, 4, 4, 4), (0.3, 0.36, 0.28, 4, 4, 4)} (respectively) with true parameter values indicated by vertical dotted line.

#### Cell Motility Rates

We analyse the effectiveness of two summary statistics which are used to estimate the motility parameters with four synthetic data sets generated with *θ* = {(0.04, 0.17, 0.08, 4, 4, 4), (0.04, 0.17, 0.08, 2, 5, 8), (0.04, 0.17, 0.08, 8, 2, 5), (0.04, 0.17, 0.08, 5, 8, 2)} where the transition rates are held constant. The first summary statistic we consider is the median position and interquartile range of each cell type on the left and right side of the scratched region; which we refer to as cell density data. The marginal posterior distributions are presented in Figure 10. We see that the estimates for cell motility are non-identifiable when cell density data is used, which is consistent with findings from Simpson et al. (2020). We believe that this may be due to interference from cells transitioning between phases and the associated difficulty in attributing the distance travelled in a phase with a single time point.

**Figure 10:**
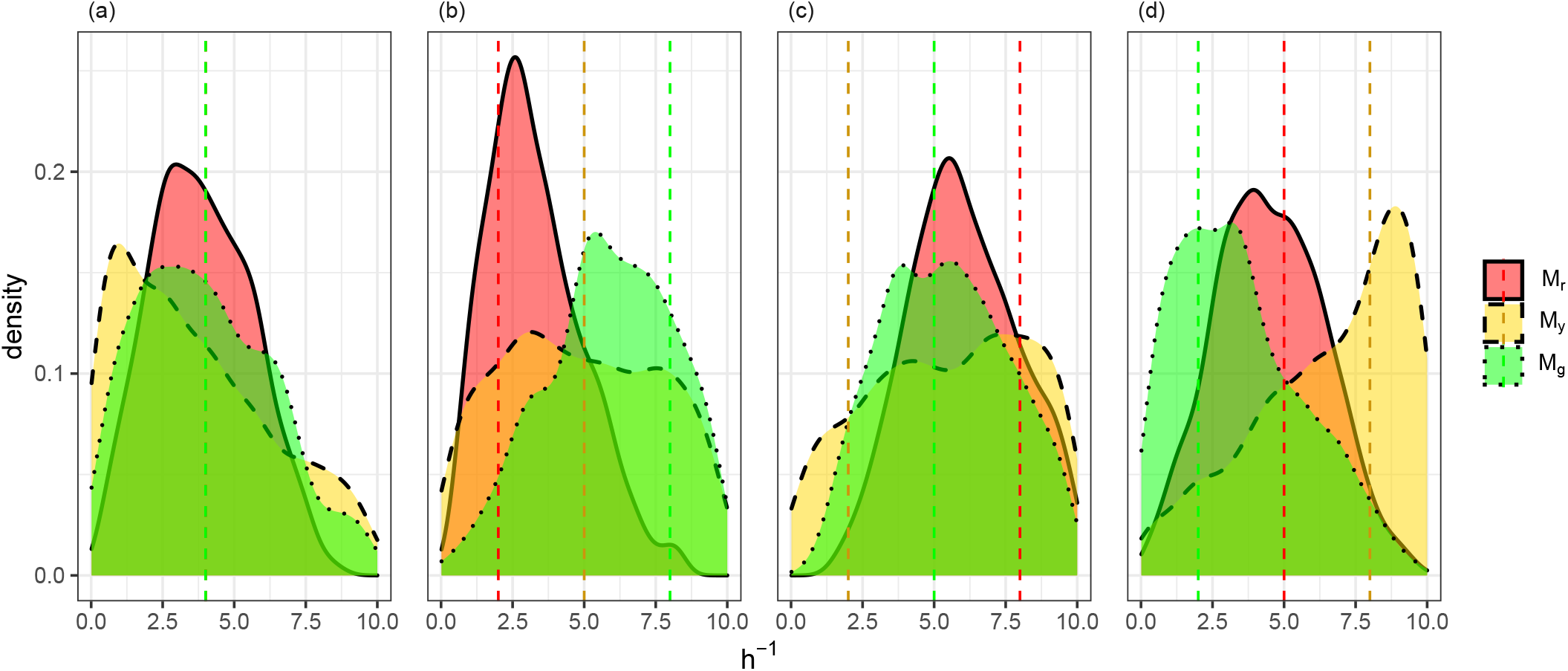
Using cell density data as summary statistics for the motility rates. (a)-(d) Posterior distributions produced using synthetic data sets generated from *θ* = {(0.04, 0.17, 0.08, 4, 4, 4), (0.04, 0.17, 0.08, 2, 5, 8), (0.04, 0.17, 0.08, 8, 2, 5), (0.04, 0.17, 0.08, 5, 8, 2)} (respectively) with true parameter values indicated by vertical dotted line.

We next consider the average distance cells travels through each cell phase of the cell cycle until the cell returns to the G1 phase or the simulation is terminated. We refer to this summary statistic as cell trajectory data and test its effectiveness with the same four synthetic data sets which were used with the cell density data with 10, 20, 30, 40 and 50 individual cell trajectories. We present the marginal posterior distributions in Figure 11. We see from the concentration of the distributions around the true value (dashed line) that cell trajectory data is highly informative about the motility rates. Furthermore, we analyse the marginal benefit of increasing the number of cells to track by 10 and find that the benefit plateaus after 20 cells. Therefore, we chose the minimally suitable number of cell trajectories which produced well defined distributions to be 20 (corresponding to Figure 11 (b,g,l,q)).

**Figure 11:**
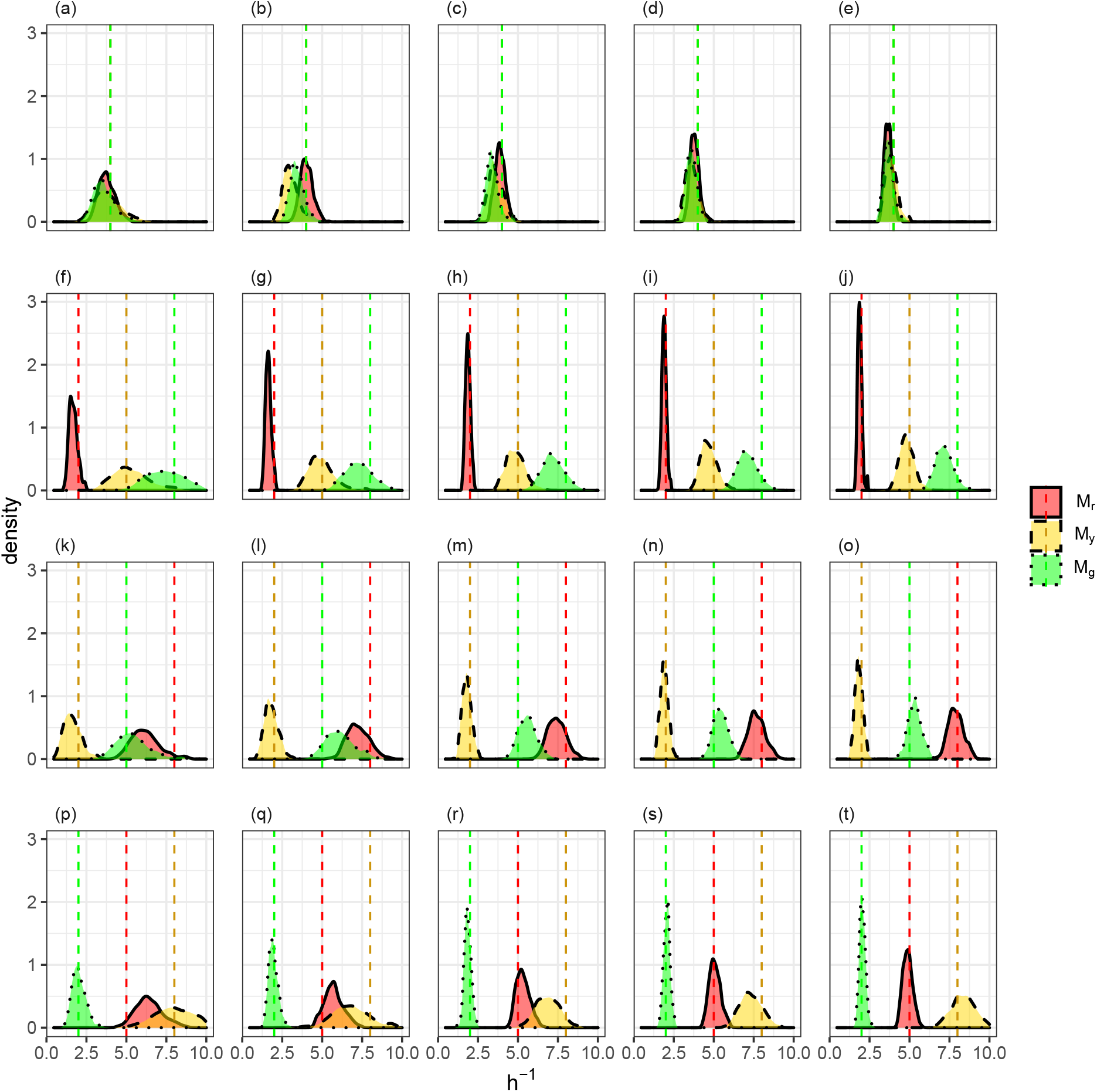
Using cell tracking data as summary statistics for the motility rates. Synthetic data sets generated from *θ* = {(0.04, 0.17, 0.08, 4, 4, 4), (0.04, 0.17, 0.08, 2, 5, 8), (0.04, 0.17, 0.08, 8, 2, 5), (0.04, 0.17, 0.08, 5, 8, 2)} are varying down the rows and number of cells tracked increases by 10 across the columns.

## References

Australian Institute of Health and Welfare. (2018). Cancer in Australia: Actual incidence data from 1982 to 2013 and mortality data from 1982 to 2014 with projections to 2017. Asia-Pacific Journal of Clinical Oncology, 14 (1), 5–15.

Beaumont, K. A., Mohana-Kumaran, N. & Haass, N. K. (2014). Modeling melanoma in vitro and in vivo. Healthcare, 2(1), 27–46.

Beaumont, M. A., Zhang, W. & Balding, D. J. (2002). Approximate Bayesian computation in population genetics. Genetics, 162(4), 2025–2035.

Blum, M. G., Nunes, M. A., Prangle, D., Sisson, S. A. et al. (2013). A comparative review of dimension reduction methods in approximate Bayesian computation. Statistical Science, 28(2), 189–208.

Cai, A. Q., Landman, K. A. & Hughes, B. D. (2007). Multi-scale modeling of a wound-healing cell migration assay. Journal of Theoretical Biology, 2f5(3), 576–594.

Codling, E. A., Plank, M. J. & Benhamou, S. (2008). Random walk models in biology. Journal of the Royal Society Interface, 5(25), 813–834.

Drovandi, C. C. & Pettitt, A. N. (2011). Estimation of parameters for macroparasite population evolution using approximate Bayesian computation. Biometrics, 67(1), 225–233.

Epanechnikov, V. A. (1969). Non-parametric estimation of a multivariate probability density. Theory of Probability & Its Applications, 14 (1), 153–158.

Ermentrout, G. B. & Edelstein-Keshet, L. (1993). Cellular automata approaches to biological modeling. Journal of Theoretical Biology, 160(1), 97–133.

Fearnhead, P. & Prangle, D. (2012). Constructing summary statistics for approximate Bayesian computation: Semi-automatic approximate Bayesian computation. Journal of the Royal Statistical Society: Series B (Statistical Methodology), 74 (3), 419–474.

Giblin, A.-V. & Thomas, J. (2007). Incidence, mortality and survival in cutaneous melanoma. Journal of Plastic, Reconstructive & Aesthetic Surgery, 60(1), 32–40.

Gillespie, D. T. (1977). Exact stochastic simulation of coupled chemical reactions. The Journal of Physical Chemistry, 81 (25), 2340–2361.

Guillemaud, T., Beaumont, M. A., Ciosi, M., Cornuet, J.-M. & Estoup, A. (2010). Inferring introduction routes of invasive species using approximate Bayesian computation on microsatellite data. Heredity, 104 (1), 88–99.

Haass, N. K., Beaumont, K. A., Hill, D. S., Anfosso, A., Mrass, P., Munoz, M. A., Kinjyo, I. & Weninger, W. (2014). Real-time cell cycle imaging during melanoma growth, invasion, and drug response. Pigment Cell & Melanoma Research, 27(5), 764–776.

Haass, N. K. & Gabrielli, B. (2017). Cell cycle-tailored targeting of metastatic melanoma: Challenges and opportunities. Experimental Dermatology, 26(7), 649–655.

Hamilton, G., Stoneking, M. & Excoffier, L. (2005). Molecular analysis reveals tighter social regulation of immigration in patrilocal populations than in matrilocal populations. Proceedings of the National Academy of Sciences, 102(21), 7476–7480.

Hastings, W. K. (1970). Monte Carlo sampling methods using Markov chains and their applications.

Ho, L. S. T., Xu, J., Crawford, F. W., Minin, V. N. & Suchard, M. A. (2018). Birth/birth-death processes and their computable transition probabilities with biological applications. Journal of Mathematical Biology, 76(4), 911–944.

Hywood, J. D., Rice, G., Pageon, S. V., Read, M. N. & Biro, M. (2021). Detection and characterization of chemotaxis without cell tracking. Journal of the Royal Society Interface, 18(176), 20200879.

Jin, W., Spoerri, L., Haass, N. K. & Simpson, M. J. (2021). Mathematical model of tumour spheroid experiments with real-time cell cycle imaging. Bulletin of Mathematical Biology, 83(5), 1–23.

Johnston, S. T., Ross, J. V., Binder, B. J., McElwain, D. L. S., Haridas, P. & Simpson, M. J. (2016). Quantifying the effect of experimental design choices for in vitro scratch assays. Journal of Theoretical Biology, 400, 19–31.

Maini, P. K., McElwain, D. L. S. & Leavesley, D. I. (2004). Traveling wave model to interpret a wound-healing cell migration assay for human peritoneal mesothelial cells. Tissue Engineering, 10(3-4), 475–482.

Marjoram, P., Molitor, J., Plagnol, V. & Tavaré, S. (2003). Markov chain Monte Carlo without likelihoods. Proceedings of the National Academy of Sciences, 100(26), 15324–15328.

Metropolis, N., Rosenbluth, A. W., Rosenbluth, M. N., Teller, A. H. & Teller, E. (1953). Equation of state calculations by fast computing machines. The Journal of Chemical Physics, 21 (6), 1087–1092.

Moler, C. & Van Loan, C. (2003). Nineteen dubious ways to compute the exponential of a matrix, twenty-five years later. SIAM Review, 45(1), 3–49.

Parkin, D. M., Bray, F., Ferlay, J. & Pisani, P. (2005). Global cancer statistics, 2002. CA: A Cancer Journal for Clinicians, 55(2), 74–108.

Pritchard, J. K., Seielstad, M. T., Perez-Lezaun, A. & Feldman, M. W. (1999). Population growth of human y chromosomes: A study of y chromosome microsatellites. Molecular Biology and Evolution, 16(12), 1791–1798.

R Core Team. (2020). R: A language and environment for statistical computing. https://www.R-project.org/

Rueden, C. T., Schindelin, J., Hiner, M. C., DeZonia, B. E., Walter, A. E., Arena, E. T. & Eliceiri, K. W. (2017). Imagej2: Imagej for the next generation of scientific image data. BMC Bioinformatics, 18(1), 1–26.

Sakaue-Sawano, A., Kurokawa, H., Morimura, T., Hanyu, A., Hama, H., Osawa, H., Kashiwagi, S., Fukami, K., Miyata, T., Miyoshi, H. et al. (2008). Visualizing spatiotemporal dynamics of multicellular cell-cycle progression. Cell, 132(3), 487–498.

Santiago-Walker, A., Li, L., Haass, N. & Herlyn, M. (2009). Melanocytes: From morphology to application. Skin Pharmacology and Physiology, 22(2), 114–121.

Savla, U., Olson, L. E. & Waters, C. M. (2004). Mathematical modeling of airway epithelial wound closure during cyclic mechanical strain. Journal of Applied Physiology, 96(2), 566–574.

Sidje, R. B. (1998). Expokit: A software package for computing matrix exponentials. ACM Transactions on Mathematical Software (TOMS), 24 (1), 130–156.

Simpson, M. J., Baker, R. E., Vittadello, S. T. & Maclaren, O. J. (2020). Practical parameter identifiability for spatio-temporal models of cell invasion. Journal of the Royal Society Interface, 17(164), 20200055.

Simpson, M. J., Jin, W., Vittadello, S. T., Tambyah, T. A., Ryan, J. M., Gunasingh, G., Haass, N. K. & McCue, S. W. (2018). Stochastic models of cell invasion with fluorescent cell cycle indicators. Physica A: Statistical Mechanics and its Applications, 510, 375–386.

Sisson, S. A., Fan, Y. & Beaumont, M. (2018). Handbook of approximate Bayesian computation. CRC Press.

Sisson, S. A., Fan, Y. & Tanaka, M. M. (2007). Sequential Monte Carlo without likelihoods. Proceedings of the National Academy of Sciences, 104 (6), 1760–1765.

Smalley, K. S., Lioni, M. & Herlyn, M. (2006). Life isn’t flat: Taking cancer biology to the next dimension. In Vitro Cellular & Developmental Biology-Animal, 42(8-9), 242–247.

Swanson, K. R. (2008). Quantifying glioma cell growth and invasion in vitro. Mathematical and Computer Modelling, 47(5-6), 638–648.

Takamizawa, K., Niu, S. & Matsuda, T. (1997). Mathematical simulation of unidirectional tissue formation: In vitro transanastomotic endothelialization model. Journal of Biomaterials Science, Polymer Edition, 8(4), 323–334.

Tavaré, S., Balding, D. J., Griffiths, R. C. & Donnelly, P. (1997). Inferring coalescence times from dna sequence data. Genetics, 145(2), 505–518.

Toni, T., Welch, D., Strelkowa, N., Ipsen, A. & Stumpf, M. P. (2009). Approximate Bayesian computation scheme for parameter inference and model selection in dynamical systems. Journal of the Royal Society Interface, 6(31), 187–202.

Treloar, K. K., Simpson, M. J., Haridas, P., Manton, K. J., Leavesley, D. I., McElwain, D. L. S. & Baker, R. E. (2013). Multiple types of data are required to identify the mechanisms influencing the spatial expansion of melanoma cell colonies. BMC Systems Biology, 7(1), 137.

Vittadello, S. T., McCue, S. W., Gunasingh, G., Haass, N. K. & Simpson, M. J. (2018). Mathematical models for cell migration with real-time cell cycle dynamics. Biophysical Journal, 114 (5), 1241–1253.

Vo, B. N., Drovandi, C. C., Pettitt, A. N. & Simpson, M. J. (2015). Quantifying uncertainty in parameter estimates for stochastic models of collective cell spreading using approximate Bayesian computation. Mathematical Biosciences, 263, 133–142.

Weyant, A., Schafer, C. & Wood-Vasey, W. M. (2013). Likelihood-free cosmological inference with type Ia supernovae: Approximate Bayesian computation for a complete treatment of uncertainty. The Astrophysical Journal, 764 (2), 116.

